## Supplementary Table S1 & S2 for "Bayesian Accounts of Perceptual Decisions in the Nonclinical Continuum of Psychosis: Greater Imprecision in Both Top-down and Bottom-up Processes"

**Supplementary Information**

***Table S1.*** *Median sensory weights significantly different from Bayesian optimal sensory weights across all conditions in both discovery and validation datasets*

| | **Discovery dataset** | | | **Validation dataset** | | |
|---|---|---|---|---|---|---|
| | Median | Optimal | p-value | Median | Optimal | p-value |
| PnLn | 0.638 | 0.465 | <2.2 x $10^{16}$ | 0.622 | 0.465 | <2.2 x $10^{16}$ |
| PnLw | 0.517 | 0.122 | <2.2 x $10^{16}$ | 0.516 | 0.122 | <2.2 x $10^{16}$ |
| PwLn | 0.828 | 0.909 | <2.2 x $10^{16}$ | 0.834 | 0.909 | <2.2 x $10^{16}$ |
| PwLw | 0.706 | 0.616 | <2.2 x $10^{16}$ | 0.725 | 0.616 | <2.2 x $10^{16}$ |

***Table S2.*** *Median subjective prior variance scores significantly different from Bayesian optimal or 'imposed' prior variance scores across all conditions in both discovery and validation datasets*

| | **Discovery dataset** | | | **Validation dataset** | | |
|---|---|---|---|---|---|---|
| | Median | Optimal | p-value | Median | Optimal | p-value |
| PnLn | 0.00105 | 0.0016 | 0.178 | 9.5 x $10^{-4}$ | 0.0016 | 0.0311 |
| PnLw | 0.00239 | 0.0016 | 6.64 x $10^{-15}$ | 2.4 x $10^{-3}$ | 0.0016 | <2.2 x $10^{16}$ |
| PwLn | 2.91 x $10^{-3}$ | 0.0185 | <2.2 x $10^{16}$ | 3.1 x $10^{-3}$ | 0.0185 | <2.2 x $10^{16}$ |
| PwLw | 0.00623 | 0.0185 | 6.77 x $10^{-16}$ | 6.4 x $10^{-3}$ | 0.0185 | <2.2 x $10^{16}$ |
